# Cyclic electron flow in *Chlamydomonas reinhardtii*

**DOI:** 10.1101/153288

**Authors:** W. J. Nawrocki, B. Bailleul, P. Cardol, F. Rappaport, F.-A. Wollman, P. Joliot

## Abstract

Cyclic electron flow (CEF), one of the major alternative electron transport pathways to the primary linear electron flow (LEF) in chloroplasts has been discovered in the middle of the last century. It is defined as a return of the reductants from the acceptor side of the Photosystem I (PSI) to the pool of its donors via the cytochrome *b*_6_*f*, and has proven essential for photosynthesis. However, despite many efforts aimed at its characterisation, the pathway and regulation of CEF remain equivocal, and its physiological significance remains to be properly defined. Here we use novel spectroscopic approaches to measure CEF in transitory conditions in the green alga *Chlamydomonas reinhardtii.* We show that CEF operates at the same maximal rate regardless of the oxygen concentration, and that the latter influences LEF, rather than CEF in vivo, which questions the recent hypotheses about the CEF supercomplex formation. We further reveal that the pathways proposed for CEF in the literature are inconsistent with the kinetic information provided by our measurements. We finally provide cues on the regulation of CEF by light.

## Introduction

Light-dependent reactions of photosynthesis couple transfer of electrons with proton translocation across the thylakoid membrane (1). Two major modes of electron transfer exist, yielding different numbers of protons pumped per electron transferred, and therefore distinctly contribute to the 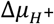 – and consequently to ATP production and luminal pH (2). Linear electron flow (LEF) - where electrons originating from water are used to reduce NADP^+^ by a work in series of both photosystems - uses 1 photon converted by each photosystem to pump 3 protons to the lumen. Cyclic electron flow (CEF) is capable of translocating two protons for each light quantum converted by PSI, but is not producing NADPH. Despite over 6 decades of research since its discovery (3), CEF mechanism, role and regulation are far from being understood, and large differences about how CEF operates are thought to exist between green algae and vascular plants, despite their obvious evolutionary relation.

CEF can be defined as a return of reducing equivalents from the PSI acceptors pool to its donor pool through the cyt. *b*_6_*f.* As vague as this definition may be, the exact pathway of the electrons back to the thylakoid membrane from the stroma is largely unknown, partly due to challenges of measuring CEF, a process with no net redox product, in vivo (4). The discovery that reduced Ferredoxin (Fd) is an efficient cofactor of cyclic photophosphorylation, even in the absence of a Fd:NADPH oxidoreductase (FNR) (5) led to propose that a putative Fd:PQ oxidoreductase catalyses CEF (6, 7). Involvement of plastoquinone was early demonstrated since cyt. *b6f* Q_O_ site inhibitors such as DBMIB and tridecyl-stigmatelin impede CEF (e.g. (8)). The pathway of CEF therefore shares at least the PQ, cyt. *b*_6_*f*, plastocyanin (PC), PSI and Fd with that of linear electron flow.

At least three separate enzymes have been proposed to fulfil the role of a [PSI electron acceptor]:PQ oxidoreductase. PGRL1 (9), a transmembrane thylakoid protein found in vascular plants as well as in green algae, has been put forward as a Fd:PQ oxidoreductase. NADH-like dehydrogenase NDH (10), a plant chloroplast homolog of mitochondrial Complex I, uses Fd or NAD(P)H to reduce plastoquinones, whereas NDA2 (11, 12) is a non-electrogenic algal NAD(P)H:PQ oxidoreductase (13), therefore they are both potentially suited to promote CEF. However, cyt. *b*_6_*f* also exhibits PQ reducing activity, and it was proposed by Mitchell as early as in the ‘70 to be a part of the original Q-cycle (14). The structural determination of the location of a *c*-type haem on the stromal side of the cyt. *b*_6_*f* (previously shown to be redox-active (15)), not present in Complex III, laid molecular foundations for the possibility of *b*_6_*f* being the FQR (16, 17). However, this role has not been proved as the mutagenesis of the *c*_i_ haem vicinity *b*_6_*f* affects the redox state of the entire Q_i_ site and thus the ensemble of electron transfer in cyt. *b*_6_*f* (Catherine de Vitry, Francis-André Wollman, unpublished).

Regulation of CEF and its rate in steady-state conditions are other important issues that shed light, less on the mechanistic aspects and pathway of the process, than on its biological role. As outlined above, CEF and LEF are in competition as they share several electron carriers. Thus, either an adjustable, heterogeneous system – where CEF and LEF actors are spatially separated and the extent of this separation can be altered - or an active, direct regulation of their kinetic constants is necessary to maintain and control these two activities.

Structuration of some photosynthetic actors is often put forward as a mean to attain heterogeneity: in algae (18) and recently in vascular plants (19), a close association between cyt. *b*_6_*f* and PSI was observed, but also FNR movement and its binding to the *b*_6_*f* (20) were hypothesised to alter CEF/LEF ratio in vivo (21, 22). A first scenario, similar to the mitochondrial multi-complex, difussionless electron tunnelling (see (23) for a review), was proposed to explain different rates of CEF in vivo, particularly those observed in anoxia (18, 24). The second scenario postulated solely that FNR would tether Fd to cyt. *b*_6_*f*, dynamically modulating the fate of Fd-carried electrons in the stroma. Kinetic regulation proposals are less abundant and consist mostly of hypotheses that dynamic redox poise of the PQ pool is a passive regulator of CEF (2, 25).

Here, we reinvestigate CEF using novel spectroscopic approaches in vivo in the model green alga *Chlamydomonas reinhardtii.* We evaluate the role of proposed CEF pathways (NDA2 and PGRL1) and compare the physiology and regulation of electron transport with those of vascular plants. A critical comment on the recent research on CEF is also provided.

## Results

### Transients of P700 oxidation reveal that CEF can operate at high rates in oxic conditions

Cyclic electron flow is notoriously challenging to measure. An access to its maximal value is however necessary to address a number of questions, notably its mechanical understanding and regulation. Until now, CEF was essayed only in steady-state conditions in the absence of PSII activity and often in non-saturating light in *Chlamydomonas:* in virtually all recent studies on CEF it is implicitly supposed that by blocking PSII, which drives LEF, one can access the rate of CEF *per se* under steady-state illumination, by measuring either the fraction of PSI reduced at a given light intensity or its photochemical rate. This hypothesis has long been proven wrong: even though partial inhibition of PSII induces CEF, a withdrawal of the PSII electron input decreases its rate (e.g. (5, 26)). We aimed at obtaining the maximal rate of CEF by inspecting *transient* conditions upon dark-to-light transition with a saturating light intensity - corresponding to over 300 electrons transferred by PSI per second (e^-^.s^-1^.PSI^-1^).

We first focused on the transients of high light-induced P700 oxidation in vivo. The redox state of the PSI special pair is easily detectable as it bleaches upon oxidation to P700^+^. The highly positive redox potential of the P700/P700^+^ couple ensures that it is the last electron carrier in the chain to become oxidised. The latter feature allows monitoring the time course of the electron transfer chain (ETC) oxidation.

The PSI donor pool in a dark-adapted state in algae is larger than in vascular plants (13) due to a more reduced plastoquinone (PQ) pool. Upon illumination in the WT, despite blocking PSII, the steady-state oxidation of thylakoid electron carriers is not attained before about 400 ms (fig. 1; see also fig. S1 for the biological variability between the experiments), during which the *number of electrons is not limiting* for CEF. However, this period is still far shorter in algae than in vascular plants (26). We therefore have relied on the PTOX2 mutant, whose PQ pool is far more reduced in darkness because it lacks the major PQ oxidase (27). Accordingly, the light-induced oxidation is more extensive in the mutant. Finally, in order to evaluate the role of PGRL1 in the CEF in *Chlamydomonas,* we also have used the PGRL1 mutant (28). As shown in the fig. 1, the oxidation of P700 in ΔPGRL1 is faster than in the WT.

**Fig. 1.**
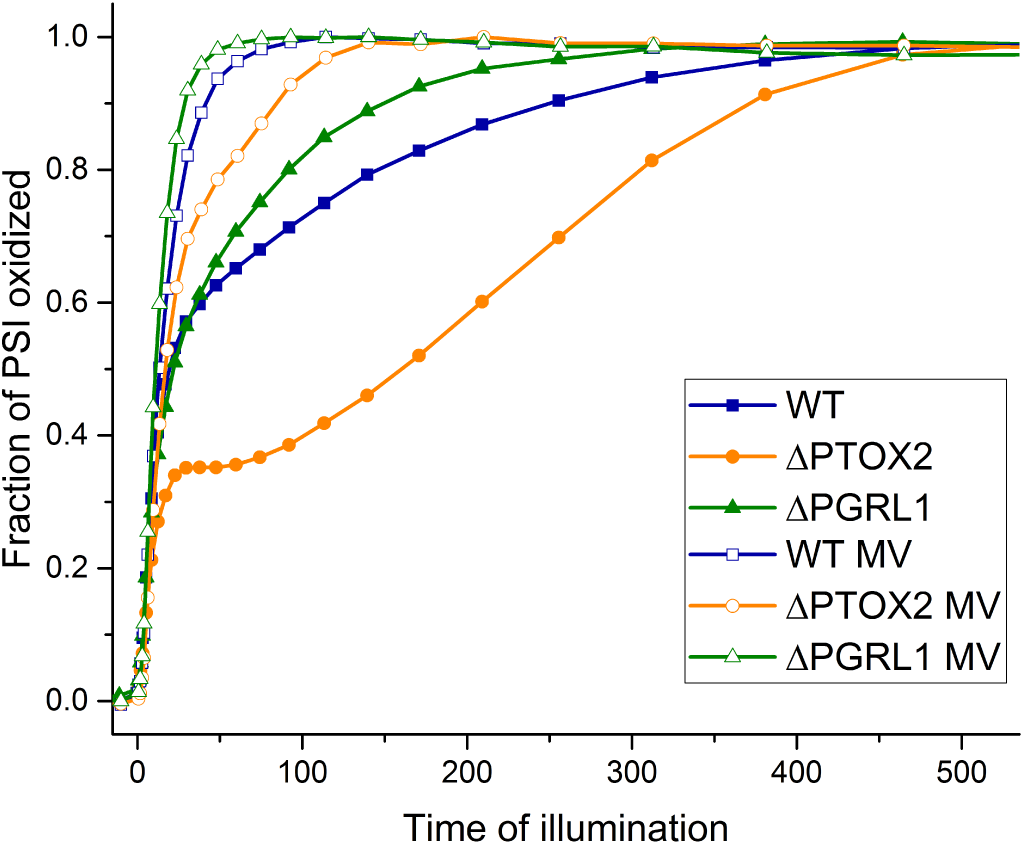
Kinetics of P700 oxidation upon dark-to-light transition in the presence of DCMU. Closed symbols – no additions. Open symbols – in the additional presence of 6 mM methyl viologen. Light intensity was set to ^~^350 e^-^.s^-1^.PSI^-1^. Averages of at least 3 independent measurements are shown.

We then sought to quantify the P700 oxidation time. Because of the marked multiphasic nature of the transients, an ambiguous fitting model would be needed to access the oxidation rates. We instead opted to use the area above the oxidation curve. We multiplied the area designated by the oxidation curve, time 0 of the illumination and the steady-state of P700 oxidation by the initial photochemical rate of PSI measured in the same conditions. This allows one to obtain the number of electrons transferred during the illumination by the PSI.

As shown in figure 4, the number of apparent PSI turnovers upon dark-to-light transition is much higher than the initial number of PSI donors present in the membrane (in the case of the WT, 40 apparent turnovers versus less than ^~^10 electron equivalents in the donor pool, e.g. (27)). This suggests that other phenomena than the sole oxidation of the pool of PSI donors contribute to the lag of P700 oxidation. Apart from CEF, also PSI charge recombination (CR) occurring when the F_A_ and F_B_ iron-sulphur clusters are reduced upon charge separation, results in a reduction of P700^+^.

We hence have used an artificial electron acceptor from PSI, methyl viologen (MV; paraquat), in order to test whether another pathway of P700^+^ reduction took place in these transitory conditions. The midpoint redox potential of about -690 mV of MV allows its di-cation form to be reduced by F_A_ and F_B_, while not oxidizing the PQ pool. As shown in fig. 1 (results quantified in the fig. 3), the oxidation of P700 took place much faster in the presence of MV, corroborating the contribution of reduction pathway(s) other than the initial donors pool.

Whereas the electron successfully transferred to the PSI acceptors pool returns to the donor pool during CEF, CR takes place when no electron acceptor is available and the non-stabilised radical pair rapidly recombines (eg. (29)). Both of these pathways would result in the same phenotype of increased lag duration. However since MV accepts electrons directly from PSI acceptors it also does not discriminate between CEF and CR.

### ECS-based measurements of photochemical rate allow discrimination between CEF and PSI charge recombination

We used a method that allows counting only stable charge separation events, i.e. giving access to the actual number of electrons transferred by PSI during dark-to-light transition and disregarding the back reactions. This method, employing the electrochromic probe of the membrane potential Δψ, DIRK (Dark-Induced Relaxation Kinetics), allows determination of the photochemical rate by comparison of the electric field formation- and dissipation rates, the latter in darkness (see (30, 31) and (32) for a review). Thanks to the detection of the rate of the Δψ decay between 1- and 4 ms after turning off the light, we exclude the contribution of rapid charge transfer across the membrane – corresponding to CR - which takes place in a sub-millisecond timescale (26, 29).

The results of these measurements are presented in figure 2. The first plotted point (at t = 1 ms) corresponds to the light-limited PSI rate (i.e. the effective light intensity) – and the initial rate of PSI in the PTOX2 mutant is higher, as shown before (24), due to its lock in state II in darkness (27) where LHCII increase PSI cross-section (33). Subsequently, the rate detected with the DIRK method in all three strains decreases until a steady-state level is attained. Whereas we observed only minor differences in rates between the WT and ΔPGRL1 in the first 60-70 ms of illumination – called the first phase hereafter -, high PSI rates are maintained for a longer period in the ΔPTOX2 during the second phase of illumination. We interpret this as a prolongation of the convoluted CEF and LEF due to an increased number of donors initially present in the membrane. Note that a normalisation of the WT curve to the maximal PSI rate in the PTOX2 mutant alleviates the differences in the rates during the first phase (see fig. S1). Additionally, one should keep in mind that the logarithmic scale used for clarity of the presentation of the initial photochemical rates is not well suited for the discrimination of differences of the areas.

**Fig. 2.**
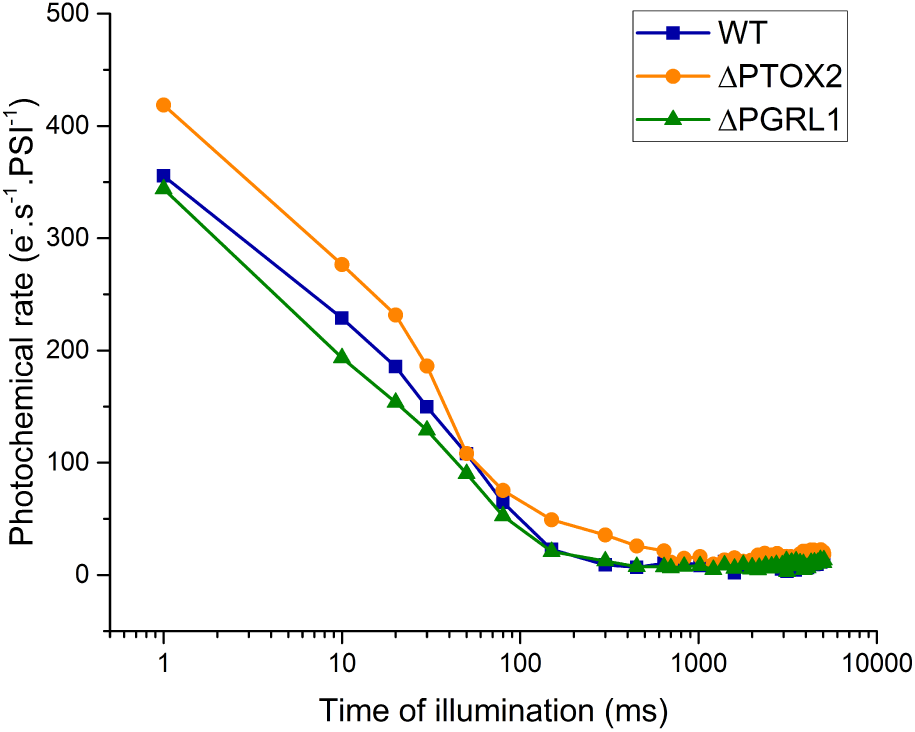
Photochemical rate of PSI during dark-to-light transition in the presence of DCMU. Each timepoint is an average of 3 to 4 biological replicates.

An integration of the curves over time until the steady-state is achieved (around 400-800 ms) yields the number of electrons transferred per PSI. Contrary to analogous integration of the P700 oxidation shown before, due to the fact that one counts only stable charge separation events with the ECS approach, only the actual electrons transferred through LEF and CEF are measured. The quantification of this integration is presented for the three strains in the fig. 3.

**Fig. 3.**
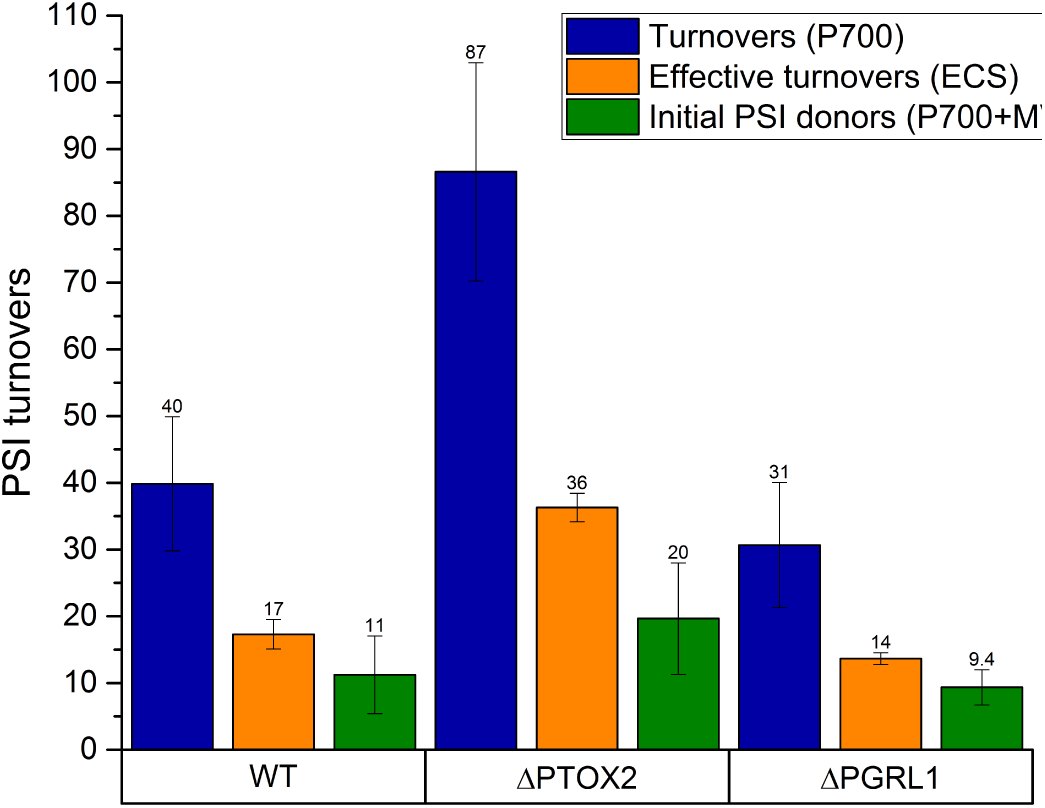
Quantification of the PSI turnovers upon dark-to-light transition in oxic conditions until a redox steady-state is attained. Blue, apparent PSI turnovers (total number of charge separations); orange, effective (stabilised) PSI turnovers; green, number of PSI turnovers in the absence of CEF and charge recombination. Means ± SDs of 3-10 independent experiments are shown.

### A combination of signal integrations yields numbers of PSI donors, electrons transferred by CEF and non-stabilised charge separations

Expectedly for the three strains, the values of the integral fall in between the number of apparent PSI turnovers and the number of initial donors in the pool. In a dark-adapted WT in oxic conditions, roughly half of the P700 oxidation area is due to CR rather than CEF+LEF. This situation changes in the PTOX mutant, where more electrons initially present in the donor pool roughly proportionally increase the number of cycles, yet drastically enhance the quantity of charge recombination. In the case of the ΔPGRL1, the pool of PSI donors is similar to the control, however both CEF and CR are affected, suggesting a general change on the PSI acceptor side.

The determination of the maximal CEF rate (V_Max_) in our conditions is challenging. Despite the use the ECS-based measurements in order to disregard back-reactions, the resulting integrals are still a convolution of CEF and LEF (i.e. LEF being the oxidation of the initial PSI donor pool without their recycling). Moreover, the presence of CR, CEF and LEF during the transient is not simultaneous. This is evidenced by the multiphasic nature of both P700 and ECS curves and by their direct comparison (fig. S1). It is visible that the first phase of oxidation, completed after ca 25 ms, is identical in the WT and ΔPTOX2, suggesting that only oxidation of the primary PSI donors occurs in this period. Next, as shown by the divergence of ECS and P700 curves, CR becomes a prominent pathway of P700^+^ reduction, followed by a CEF phase about 100 ms after the onset of illumination. Then, CEF operates at about 75 e^-^.s^-1^.PSI^-1^. This result shows that in oxic conditions in *Chlamydomonas,* CEF is able to work at high rates, identical to the maximal values reported to date in anoxia and assigned to be a result of CEF regulation. The CEF phase lasts significantly longer in the ΔPTOX2 mutant suggesting that the lower number of electrons causes its rapid decay in the WT. However, due to the extremely short duration of the oxidation phase (especially compared to the parallel situation in vascular plants (26)), the accuracy of the CEF V_Max_ estimation is low and may be underestimated because of this electron limitation. We have thus employed hypoxia in a bid to, at a time, increase the CEF duration and investigate its response to various oxygen concentrations, and also to verify whether the *maximal rate* of CEF increases in reducing conditions.

### Varying the oxygen tension allows prolongation of the rapid transitory CEF

It has been previously shown that the lag in P700 oxidation increases in lower oxygen concentration conditions, with the extreme case being anoxia where initially none of the centres can be oxidized (e.g. (34)). Here we present an extension of the new ECS-based approach to a range of oxygen concentrations (fig. 4).

**Fig. 4.**
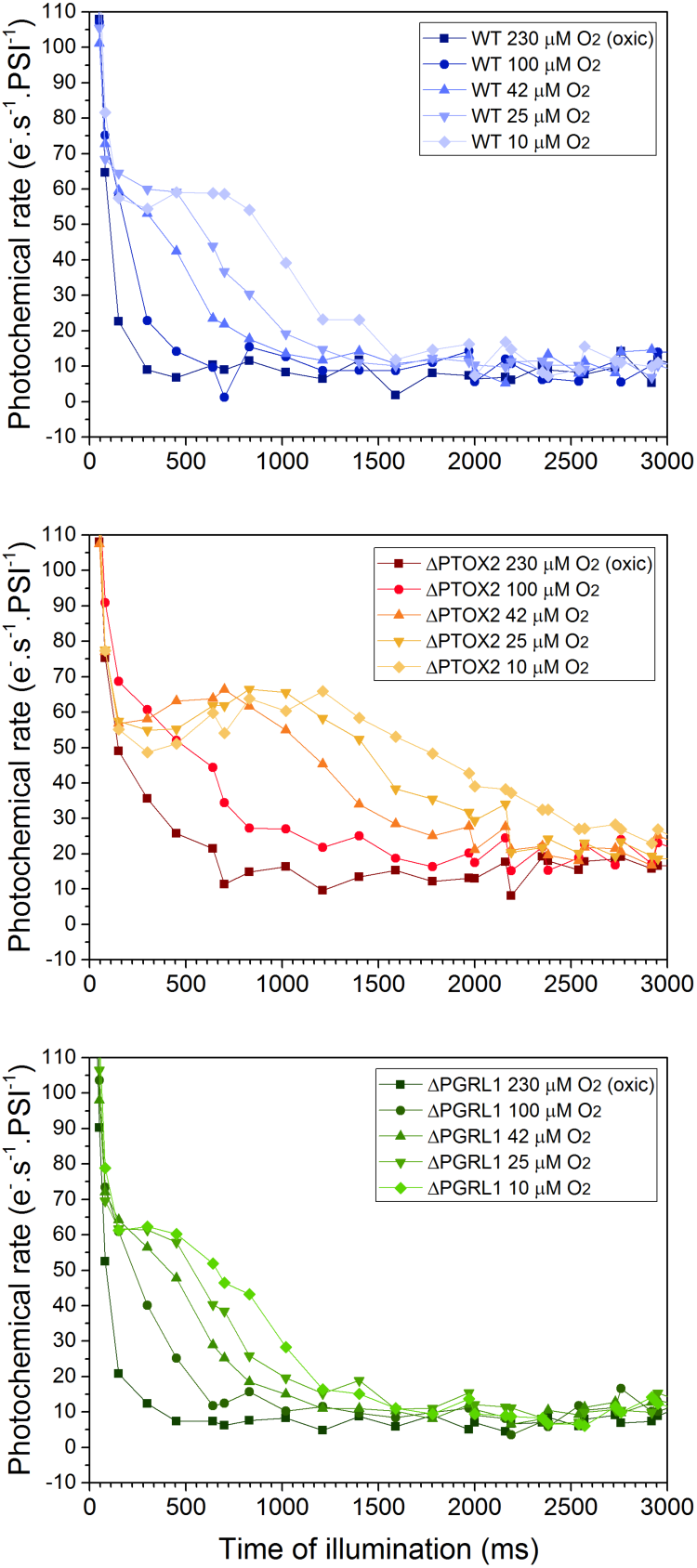
Photochemical rate of PSI during dark-to-light transition in the presence of DCMU as a function of oxygen concentration. A zoom on the second phase is shown (see text for details). Each timepoint is an average of 3 biological replicates.

Upon decreasing oxygen concentrations, a progressive increase of the duration of the elevated PSI photochemical rates was observed during the second phase of illumination in all strains. We interpret this phase as pure CEF at a rate close to its maximum capacity, in this case about 70 e^-^.s^- 1^.PSI^-1^, consistent with the above estimations in oxic conditions. Whereas in the WT a steady-state was attained at around 200-400 ms in oxic conditions (i.e. ^~^230 μM O_2_), it took up to 1.5 seconds to achieve it in the extreme case of 10 μM O_2_. While the maximal rate of PSI does not change during the second phase, it is the duration of the CEF period that increases. This is corroborated by the results with the ΔPTOX2 mutant, where the high rates even in oxic conditions are prolonged with regards to the WT. Importantly, in the ΔPGRL1 mutant the overall kinetics and absolute values are nearly identical to those in hypoxic WT, with a slightly shorter CEF phase, similarly to the situation in oxia (Figs 2 and S1). This again – similarly to oxic conditions – suggests that some unknown redox differences exist in the ΔPGRL1 on the PSI acceptor side, and not that the CEF mechanism itself is affected. The slight shortening of the duration of the oxidation phase in ΔPGRL1 may explain why previously the mutant has been thought to be unable to perform CEF in anoxia (35).

Finally, an increase in the maximal, light-limited PSI rate is shown to occur with a decrease of oxygen concentration (fig. S7). This corroborates previous results showing that the PSI antenna size increases in anoxia, upon State Transitions (33). Little differences are observed in the ΔPTOX2 mutant, in line with its lock in state II in darkness (27).

One needs to distinguish between two possible reasons for the increase in the area of P700 oxidation in hypoxia, that is an increase in the number of PSI donors or a decrease of the electron outflux to PSI acceptors. In the first case, chlororespiration may be affected as the concentration of oxygen, the substrate of PTOX, decreases. On the other hand, PSI acceptor side limitation can explain the increased area as both the Mehler reaction and the flavo-diiron proteins lack their substrate. In addition the increased reduction in the stroma should decrease the activity of the primary electron sink, CBB cycle. We have probed, using fluorescence induction, which of these hypotheses can explain the lag in hypoxia (Figs S3 and S4). Whereas the very first phase (below 200-400 ms) represents a probe of the redox state of the initial PSI acceptors and then donor side pools, the second phase up to 5 seconds exhibits a quasi steady-state of electron flow. We observed a modest increase in the PSI donor and acceptor pool reduction as evidenced by the F0 level and initial kinetics of the fluorescence in light. This reduction explains the state I to state II transition as evidenced by the PSI antenna size changes shown above (Fig. S5). However, after an increase in fluorescence, for 250 ms, a decrease in F_Stat_ is observed in high oxygen concentration, which is absent in hypoxia. We note here that in order to achieve a good temporal resolution, we used a slightly lower actinic light intensity for this experiment as compared to the previous ones. Furthermore, DCMU had been consistently used, apart from these fluorescence measurements, the result being that the oxygen-sensitive oxidation is delayed with regards to the CEF decrease presented in the figure 2, the latter occurring about 100 ms after the beginning of illumination in oxic conditions. Nonetheless, these results show that primarily the decrease of an electron leak, and not the initial electron supply or redox steady-state before illumination, is behind the increase of the lag of P700 oxidation in hypoxia.

Taken together, these results suggest that the progressive increase in CEF duration in hypoxia is due to a decrease of the leak of electrons toward oxygen. Therefore differences in LEF, and not a contrived regulation or structural changes increasing CEF V_Max_ alter the CEF duration in hypoxia, which was before incorrectly interpreted as a change of its rate.

### Preillumination studies argue for CEF regulation in *Chlamydomonas*

We then focused on the regulation of CEF. We have studied the effect of a preillumination in the presence of DCMU, and different dark periods that followed, on the P700 oxidation. The area was quantified in the same manner as in the experiments above, and the results are shown in the fig. 5. Initially, the area below P700 oxidation curve is small, due to the preillumination that oxidizes the pool of PSI donors. As the time in darkness increases, a reduction of the PSI donor pool occurs. Then, after a maximal value is attained, the areas shrink until a steady-state is achieved after 1 to 3 minutes of darkness (last point on the x axis in the fig. 5). This decrease is monotonous. The shape of an overshoot of the kinetics suggests that at least two competing processes with different rate constants contribute to the observed phenomenon. This is in line with our previous observations, where in the dark-adapted sample (i) the initial state of the PSI donor pool, (ii) CEF and (iii) charge recombination all contribute to the area increase.

**Fig. 5.**
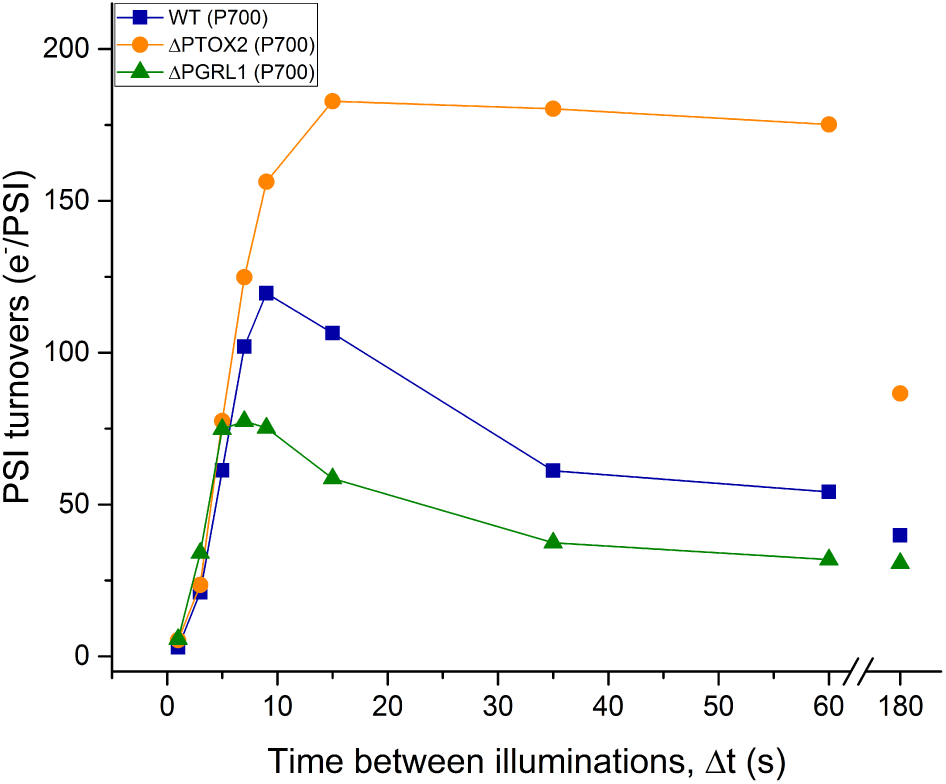
Evolution of the apparent number of PSI turnovers before attaining a redox steady-state in the presence of DCMU as a function of time in darkness following a preillumination. Each timepoint is an average of at least 3 biological replicates.

Importantly, the areas vary between WT and the mutants. Whereas the first phase of the kinetics, corresponding to the reduction of the oxidized PSI donor pool is similar in all the strains, the second phase (from about 10 seconds to the dark-adapted state) is strikingly slower in the ΔPTOX2 mutant, and the amplitude reaches 200 apparent PSI turnovers upon PSI oxidation. On the other hand, whereas the kinetics of the second phase is similar in the WT and the ΔPGRL1 mutant, the amplitude is lower for each time point in the latter.

We then used the ECS-based method to investigate whether CR can explain the large apparent number of turnovers in the preilluminated samples. The results (compare fig. 6 with fig. 5) suggest that indeed the contribution of CR to the P700 area during dark-to-light transition is high. This is particularly the case for a transient period between 5 and ^~^30 seconds in the WT and the ΔPGRL1 mutant; in the case of ΔPTOX2, CR is present for a far longer period. However, in all cases the shape of the curve is similar – and consists of a first phase where CEF+LEF increase, and a slower second phase of apparent oxidation until steady-state – suggesting that the regulation of CEF is highly similar in those strains.

**Fig. 6.**
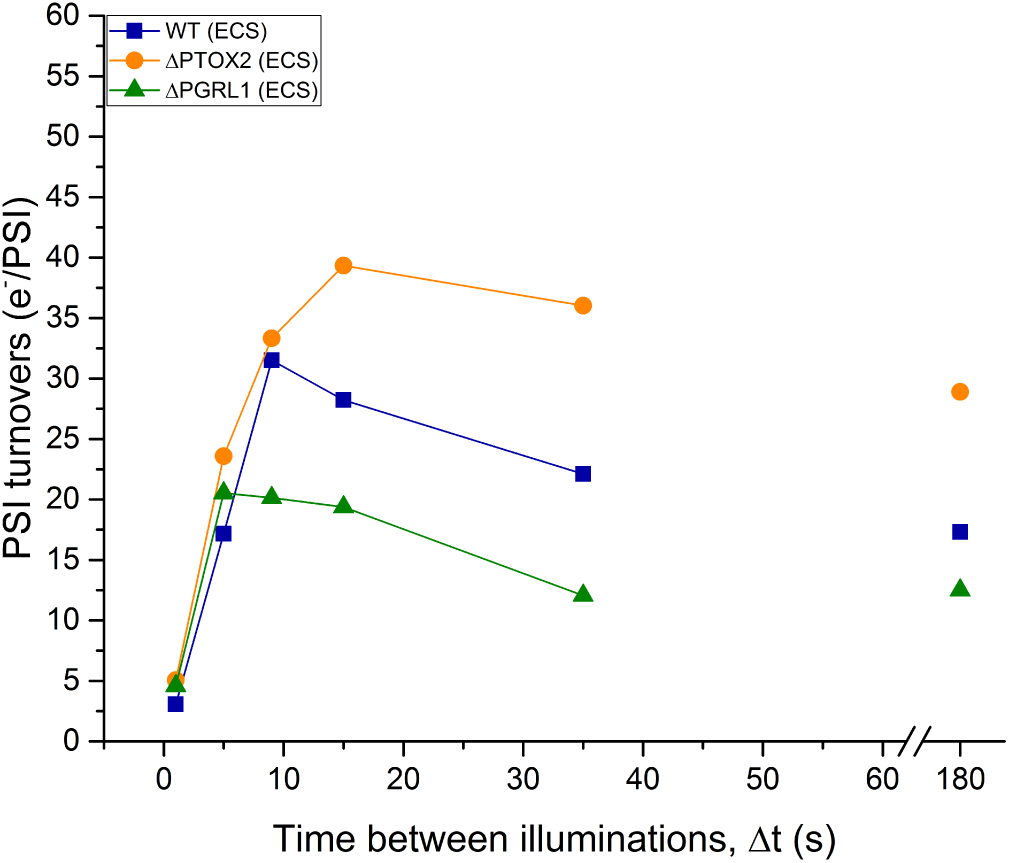
Evolution of the number of stable PSI turnovers before attaining a redox steady-state as a function of time in darkness following a preillumination. Each timepoint is an average of at least 3 biological replicates and is a result of an integration of the area below the curve analogous to those presented in the fig. 2.

Finally, we have quantified the area below preilluminated P700 oxidation curves in the presence of methyl viologen. The results, presented in fig 7., consist of two phases in all strains. We interpret the first phase of increase in the number of PSI turnovers as a reduction of the pre-oxidized PSI donor pool. After about 10 seconds in darkness, this process is finished and a steady-state is achieved. Whereas the latter is virtually identical in the WT and ΔPGRL1 mutant, the number of PSI turnovers needed to oxidize PSI is higher in the ΔPTOX2 mutant, consistent with an increased amount of PSI donors in darkness.

**Fig. 7.**
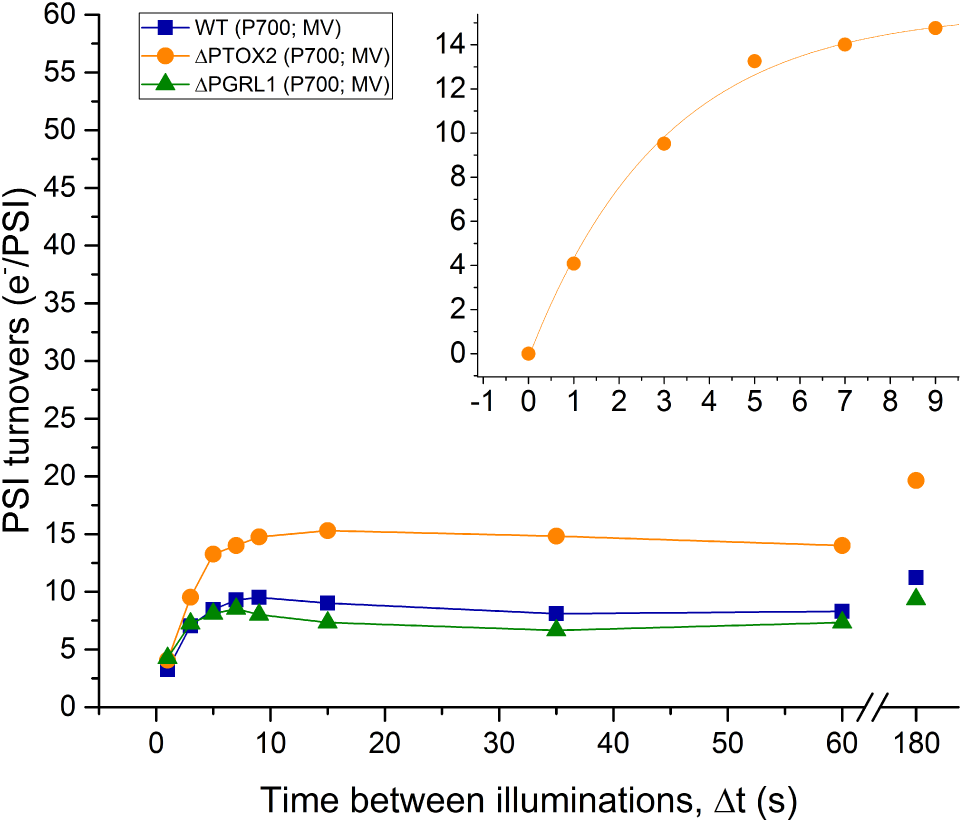
Evolution of the number of PSI turnovers before attaining a redox steady-state in the absence of CEF and charge recombinations as a function of time in darkness following a preillumination. Each timepoint is an average of at least 3 biological replicates.

The rise in the area below the P700 oxidation curve has a logarithmic shape, suggesting that a single process, or two processes with similar rate constants contribute to the observed increase. This is consistent with the model of chlororespiration presented before (27), where two activities, reducing and oxidizing, dynamically balance the redox equilibrium of the PQ pool. We conclude that in the presence of MV the number of PSI electron donors can be obtained. This number is shown to be predominantly determined by chlororespiration, which regulates the redox state of the PQ pool, and to be similar in the WT and ΔPGRL1 mutant.

Last, an analysis of the kinetics of the PSI donor pool reduction in the presence of MV in the PTOX2 mutant allows a cross-validation of our method with previous, fluorescence-based approach in a *b*_6_*f* -lacking mutant, aimed at a determination of the maximal rate of NDA2. A mono-exponential function was used to fit the reduction of the PQ pool (fig. 7, inset) in the absence of the major PQ oxidase. We could estimate the rate of NDA2 to be slightly higher than 4 e^-^.s^- 1^.PSI^-1^, a value comparable to the one obtained beforehand (2 - 2.4 e^-^.s^- 1^.PSI^-1^; (27)).

## Discussion

CEF characterisation is hampered by its intrinsic convolution with linear electron flow and the absence of measurable net products. There is a two-way relation between LEF and CEF, where the former on one hand supplies CEF with electrons by which it *increases* CEF rate, yet it also directly competes with CEF for several intermediate electron carriers, therefore limiting it. We approached this conundrum by focusing on a transient situation, where although LEF is blocked, the initial pool of electron donors for CEF is relatively large. This strategy proved fruitful because it allowed monitoring CEF that was temporarily not electron-limited and operating at high rates that we hypothesize are close to the CEF V_Max_.

While investigation of P700 oxidation and ECS-based PSI charge separation rates allowed determination of the apparent- and actual number of PSI turnovers until the steady state was reached, addition of an artificial electron acceptor enabled quantification of the initial size of PSI donor pool. The number of turnovers in the presence of MV in all strains is close to the data from the literature (compare with (27)). Conversely, the rate of NDA2 estimated from the MV experiments is higher than reported previously – we hypothesize that some residual CEF/CR, not completely abolished by the competitive inhibitor MV, still operates in these conditions.

### Revisiting the pathway of CEF

Long-lasting efforts to isolate mutants of CEF contributed to our understanding of the molecular bases of this process. Almost exclusively, two CEF routes are considered in the literature to be responsible for the reduction of PQ by PSI donors: the NDH/NDA2 and PGR5/PGRL1-dependent pathways (eg (36)). The components of the first pathway are homologous to the respiratory NAD(P)H:PQ oxidoreductases: the NDH to complex I, and NDA2 to NDI1. Even though NDH has been proposed to use Fd as substrate (10) and pumps additional protons per electron transferred (13), its role in CEF is doubtful from many perspectives. NDH is highly substoichiometric with regards to PSI (1:100 ratio), therefore to sustain observed rates of CEF in plants of ^~^130 e^-^.s^- 1^.PSI^-1^ its rate would necessarily need to exceed 10^4^ e^-^.s^-1^ (21, 26). The proposed structuration of PSI-NDH (37) fails to explain such high hypothetical rates in a diffusion-limited complex since cyt. *b*_6_*f* was not shown to be in close interaction with these proteins. In addition it would imply that both Fd and Pc need to operate as intra-complex electron carriers. Last, the rate of PQ oxidation at the Q_O_ site would set the kinetic limitation to a few hundred e^-^.s^-1^ which is far too low. Finally, experimental data shows that the NDH does not contribute to q_E_ (pH-dependent NPQ) – and therefore to the proton gradient – in plants (38), and its rate was measured at a fraction of an e^-^.s^- 1^.PSI^-1^ in vivo (39). Similarly, NDA2 - which is the algal analogue of NDH – is at its 2.5 e^-^.s^- 1^.PSI^-1^ (13) far too sluggish to sustain CEF in the order of 70 e^-^.s^-1^.PSI^-1^ that we observe. Finally, only differences in photosynthesis observed in the NDA2 mutant compared to the wild-type regarded non-photochemical, slow PQ reduction (12) or when the competing PQ-reducing PSII activity was severely affected (40).

The second most coveted CEF route is thought to be PGR5/PGRL1-dependent. The two molecular actors of this pathway were discovered in mutagenized plants exhibiting distinctly low steady-state q_E_, and thus ΔpH across the thylakoid membrane, leading to their name (proton gradient regulation 5 and pgr5-like 1, respectively). However, a considerable amount of data argues against the direct involvement of this pathway in CEF. Whereas the PGR5 has already been shown to only *regulate* CEF (41), PGRL1 is still thought to fulfil the role of the elusive FQR (9) directly reducing quinones. This is incompatible with several observations: the PGRL1 is also sub-stoichiometric with regards to the electron transfer chain, with about 0.3 proteins per PSI, or 0.15.PSI^-1^ if it dimerises in vivo (9). Most of all, our measurements presented above show that no differences in the V_Max_ of CEF, no modification of the redox state of the PQ pool nor alterations in the CEF regulation are observed in the mutant, all of which would be expected if PGRL1 was an FQR. Rather than that, the PSI acceptor side is generally affected with both CEF and CR probability being lower than in the WT, a feat suggesting a role of PGRL1, similar to that of PGR5, in regulating the fate of PSI donors or the redox state of the stroma, but not directly CEF. This is finally consistent with the observation that an overexpression of Flvs in a ΔPGR5-background plants restores their WT phenotype (42).

Taken together, these considerations strongly support the third possible route for the return of electrons to the PSI donor side – the direct PQ reduction by cyt. *b*_6_*f*. Originally proposed by Mitchell and reiterated afterwards (14, 26), the classical Q-cycle consists of a double reduction of the PQ at the Q_i_ site. Whereas one electron is borne from the *b*_h_ haem (following a quinol oxidation at the Q_o_ site), the second could be coming from stromal donor such as Fd, possibly through the redox-active *c*_i_ haem (15, 16). Accordingly, FNR was shown to bind to the *b*_6_*f* and regulate CEF (20, 21). It could therefore act as a tether for the Fd in the vicinity of stromal haems of the *b*_6_*f*, or even be involved in the electron transfer directly (Fd in FD:FNR complex has a redox potential of -400 to -500 mV, FNR -350 mV (43); *b*_h_ haem is - 30 mV, *c*_i_ +100 mV but in vivo the redox difference of the two is much smaller and in dark-adapted algae the *c*_i_ haem is more reduced than the *b*_h_ haem; see (44) for a discussion). If this is the pathway of CEF in vivo, the only electron carrier not shared between the two major modes of electron transfer would be the *c*_i_ haem, making it a primary target for the approaches testing this hypothesis. However, alterations of the *c*_i_ haem strongly influence the neighbouring *b*_h_ haem, yielding this method inapplicable. The fact that in practice zero carriers are specific to CEF could explain the long-standing difficulties in obtaining null mutants of this route.

### What regulates CEF in hypoxia?

The short period when CEF operates at high rates in our experiments in oxic conditions (from ^~^80 to ^~^100 ms after the beginning of illumination) led us to attempt to limit the electron leak from CEF toward downstream photosynthetic reactions. It has long been known that CEF is enhanced when oxygen concentration decreases (e.g. (24, 34, 45)). We were able to increase the duration of rapid CEF in hypoxic conditions for up to 2 seconds. Importantly, thanks to the progressive, respiration-mediated oxygen depletion in the sample, this prolongation is continuous. All together these results allowed us to conclude that the previously observed increased rates of CEF in anoxia (18, 24, 34, 35, 45, 46) did not occur because of differences in structuration (like PSI-*b*_6_*f* supercomplex formation or state transitions) or CEF regulation (redox regulation), but simply because of the suppression of electron leak competing with CEF. We were then able to exclude that initial redox state of the PQ pool or PSI acceptors was the reason for a lag in PSI photochemistry in hypoxia, but rather that it was the suppression of oxygen-reducing reactions which caused CEF enhancement. The recent discovery of flavo-diiron proteins (Flvs) in Chlamydomonas (47, 48) is consistent with these observations – especially given that Flvs are absent in angiosperms where the PSI oxidation lag in the presence of DCMU is lengthy (26). We exclude the involvement of the Mehler reaction - a direct oxygen reduction by the F_A_/F_B_ ISCs - because it is not sensitive to oxygen in our range of concentrations. The oxygen titration presented above is, to our knowledge, the first demonstration that the affinity of Flv for oxygen has a fairly low apparent KM of ^~^50 μM in vivo.

### Regulation of CEF/LEF partitioning

CEF competes with LEF for at least two electron carriers: oxidized PQ and reduced Fd. The regulation of the relative activities of these two processes in vivo will necessarily need to alter the rate at which those transporters are reduced and/or oxidized. But is it necessary to regulate both PQ and Fd sharing? We would like to argue here that a control of the fate of the electrons in the stroma is sufficient for a modification of CEF/LEF partitioning, and that PQ redox state is not influencing CEF. It was shown previously, using sub-saturating laser flashes in anoxic conditions, that a quinone readily produced (oxidized) at the Q_o_ site has an absolute priority to shuttle to the Q_i_ pocket of the very same complex (49), rather than escape from the same complex and diffusionally explore Q_i_ sites in its vicinity.

We propose that the inverse happens in vivo, that is nascent quinols - reduced at the Q_i_ site – have a priority for the oxidation at the adjacent Q_o_ site of the same cyt. *b*_6_*f*, i.e. without entering the “PQ pool”; that constitutes a heterogeneity within the distribution of the electron transfer carriers. This heterogeneity would in effect translate to two different routes to the Q_o_ site, one for “LEF quinols” from outside of the complex, and the other one from “CEF quinols” from the Q_i_ site, thanks to the latter being deeper inside the cavity in the *b*_6_*f* dimer. Such mechanism effectively uncouples the redox state of the PQ pool form the CEF rate - quinone reduction at the Q_i_ is *independent* from the overall state of the pool and thus from competing reduction of the PQ by PSII. Such a condition has been previously observed: in transitory situation when the PQ pool was almost entirely reduced by strong illumination, CEF in *Chlamydomonas* prevailed over LEF reaching rates close to the V_Max_ (50). This stands in disagreement with previous proposals (2, 25) that the highest efficiency of CEF is achieved when 50% of the PQ pool is oxidized, and 50% in reduced state.

Consequently, competition between CEF and LEF for reduced Fd is in our view the hub where those two pathways necessitate regulatory actions. As outlined before, Fd^red^ can be readily oxidized by the CBB cycle and apart from plants, by flavodiiron proteins (47). Such oxidation was shown here to be decreased in low oxygen concentrations, and an increase of the *duration* of CEF also observed in Flv mutant of moss (51), yet it was interpreted as a compensatory increase in CEF rather than a decrease of the electron leak.

Regulation between CEF and LEF in this case could be achieved as proposed before, by increasing the probability that ferredoxin stays in proximity of the stromal side of the cyt. *b*_6_*f* (21), through an anchoring by the *b*_6_*f* -bound FNR. Recently, FNR has been shown to be a target of the redox-sensitive STN kinase (52), but functional and biochemical data regarding the regulation of Fd-FNR- *b*_6_*f* binding is scarce at the moment. Fd is at the very crossroads of photosynthetic electron transfer, donating electrons not only to CEF and LEF, but also participating, among others, in nitrite and sulphide reduction. Furthermore, multiple Fd isoforms, with varying redox potentials and concentrations were shown to exist, and moreover they interact with multiple isoforms of FNR, some of which are soluble and some membrane-bound. It is obvious that a strict regulation of this hub is necessary for photosynthesis and the redox state of the entire stroma, making it easy to imagine that CEF is also governed at this level.

### What is the V_Max_ of CEF in Chlamydomonas?

It is important to obtain the sought-after value of the maximal rate of CEF, and the conditions where it occurs. In vascular plants, low-energy chlorophylls in LHCa allow partially specific PSI excitation by far-red light, in which case the initial, small electrons leak to downstream reactions is compensated by PSII activity without compromising PSI activity. In these conditions, CEF of up to 130 e^-^.s^-1^.PSI^-1^ has been observed (26). In *Chlamydomonas,* the spectral difference between the Photosystems is too small for this method to be employed (53, 54), but on the other hand the initial number of PSI donors in dark-adapted sample is higher due to differences in chlororespiration (13). This allowed us to study CEF in the absence of PSII activity, in the transitory period before a redox steady-state was achieved. In this case, CEF and LEF are intermingled, yet qualitative information about CEF rate can still be estimated – in the WT in oxic conditions the effective rate of CEF was in the order of 70 e-.s^-1^.PSI^-1^. Furthermore, decreasing the electron leaks downstream PSI allowed a 10-fold prolongation of CEF in order to make sure that the elevated rates of CEF observed in oxic conditions are not an artefact of our measurements. This rate could still be slightly underestimated as in the light we used, corresponding to 350 e^-^.s^-1^.PSI^-1^, CEF in vascular plants is not yet saturated (P. Joliot, unpublished). We conclude that transitory conditions where the PQ pool is largely – but not entirely – reduced, together with a decreased electrons leak to downstream reactions and a careful use of the method that allows observation of stable PSI charge separations in strong light are needed to fulfil the requirements of an experimental setup for CEF V_Max_ measurements in *Chlamydomonas*.

## Acknowledgements, statements

W.J.N was supported by French Ministry of Education. P.C is a Senior Research Associate from Belgian F.R.S.-FNRS. The authors acknowledge funding from the ERC (consolidator grant BEAL 682580), French state (Labex DYNAMO ANR-11-LABX-0011-01), CNRS and UPMC.

## Materials and methods

WT (CC124), ΔPTOX2 (27), and ΔPGRL1 (28) were grown at 25°C in continuous 10 μE.m^-2^.s^-1^ light in Tris-acetate-phosphate media. Prior to each experiment, the cells from mid-log phase were spun down and resuspended in 10% (w/w) Ficoll-Minimum medium (without reduced carbon source), then dark-adapted in open Erlenmeyer flasks and shaken for at least 1 hour. Apart from the fluorescence measurements (Figs S3, S4 only), DCMU (10 μM) and hydroxylamine (10 mM) were systematically added in order to block PSII.

All the spectroscopic measurements were done using a JTS-10 spectrophotometer and orange actinic LEDs peaking at 635 nm. Single turnover flashes used to normalize the ECS values were obtained through a dye laser (DCM, Exciton dye laser) pumped by a frequency-doubled Nd:YAG Laser. P700 redox state was measured at 705 nm (white LEDs passing through an interference filter with 10 nm full width at half maximum) and subtraction of a corresponding measurement at 730 nm (10 nm FWHM) to remove PC-borne signals was made. ECS was monitored at 520 nm (white LEDs passing through an interference filter with 10 nm FWHM). Actinic light was cut off from the detector by long-pass Schott RG695 glass filters and BG39 Schott filters, respectively. The light in all the experiments corresponded to over 300 PSI turnovers per second.

The oxygen measurements were performed using contactless Sensor Spots (PyroScience) stuck inside the measuring cuvette and simultaneously with the JTS-10 measurements. The sensors were calibrated before the experiments with Ficoll-Minimum medium bubbled with air for oxic condition, and with DTT added to the medium in closed cuvette for anoxia reference.

Detectable electric field across the thylakoid membrane is formed and dissipated by photosynthetic complexes in the light. Upon cessation of illumination the photochemical rate is instantly cancelled, yet on a timescale of several milliseconds the light-independent potential formation is unchanged, allowing an absolute quantification of the PSI photochemistry in the case PSII is inhibited. While charge recombination is electrogenic and contributes to the ECS signal as the electrons move from the stromal to the luminal part of the membrane, it decays rapidly in PSI (29). By disregarding the ECS signals before 1 ms of darkness – when the vast majority of back-reactions takes place - one is able to only measure the stable rate of charge separations, i.e. ‘forward’ PSI photochemistry.

The photochemical rate of PSI along the whole period of P700 oxidation using the ECS-based approach was measured as follows. Each timepoint corresponds to a single-turnover flash-normalised value of a difference between ECS slope in the light *minus* dark. The slopes were obtained through a linear fit of 8 1-ms-spaced detection points in the presence of actinic light, and 4 1-ms-spaced points after the light was turned off. Importantly, the first detection point in darkness was made after 1 ms to avoid the contribution of charge recombination. Because a short, 5-ms period of darkness was necessary for the measurement for each timepoint, the measurement was split into two, where either timepoints number *n, n+2*, *n+4*, etc. or *n+1*, *n+3*, *n+5*, etc. were recorded. This avoided an artificial decrease in the perceived light at the beginning of the measurement, where the points are closely spaced. Finally, a baseline consisting of a measurement without the actinic light was subtracted from every set of data. Each of the points is a result of 5-8 technical replicates (of same batch of algae) of the ECS measurement and averaged over 3 biological replicates.

## Supplementary materials

**Fig. S1.**
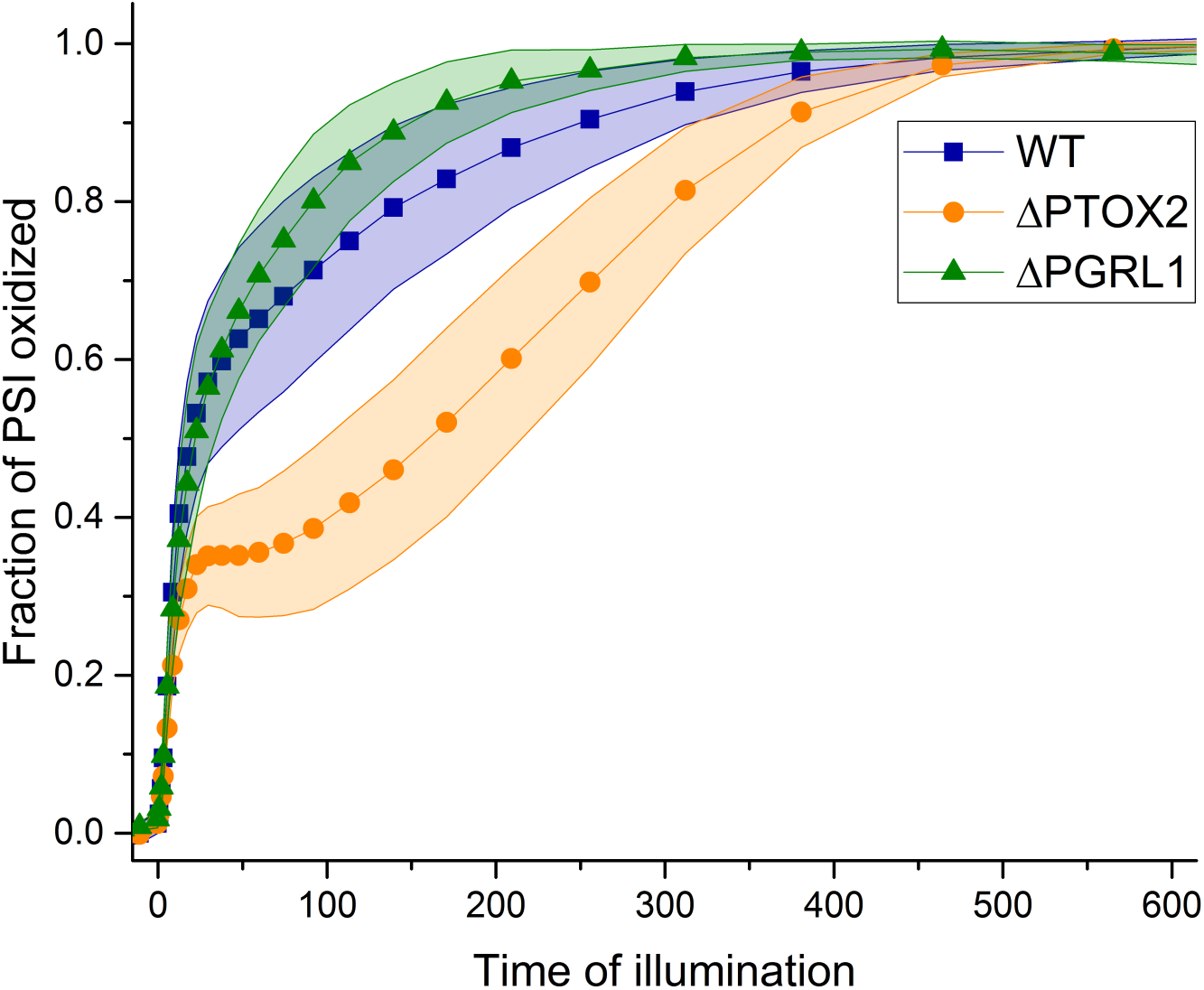
Biological variability in the P700 oxidation kinetics in dark-adapted samples. Shaded areas correspond to the standard deviation from the mean (closed symbols) of at least 3 independent measurements.

**Fig. S2.**
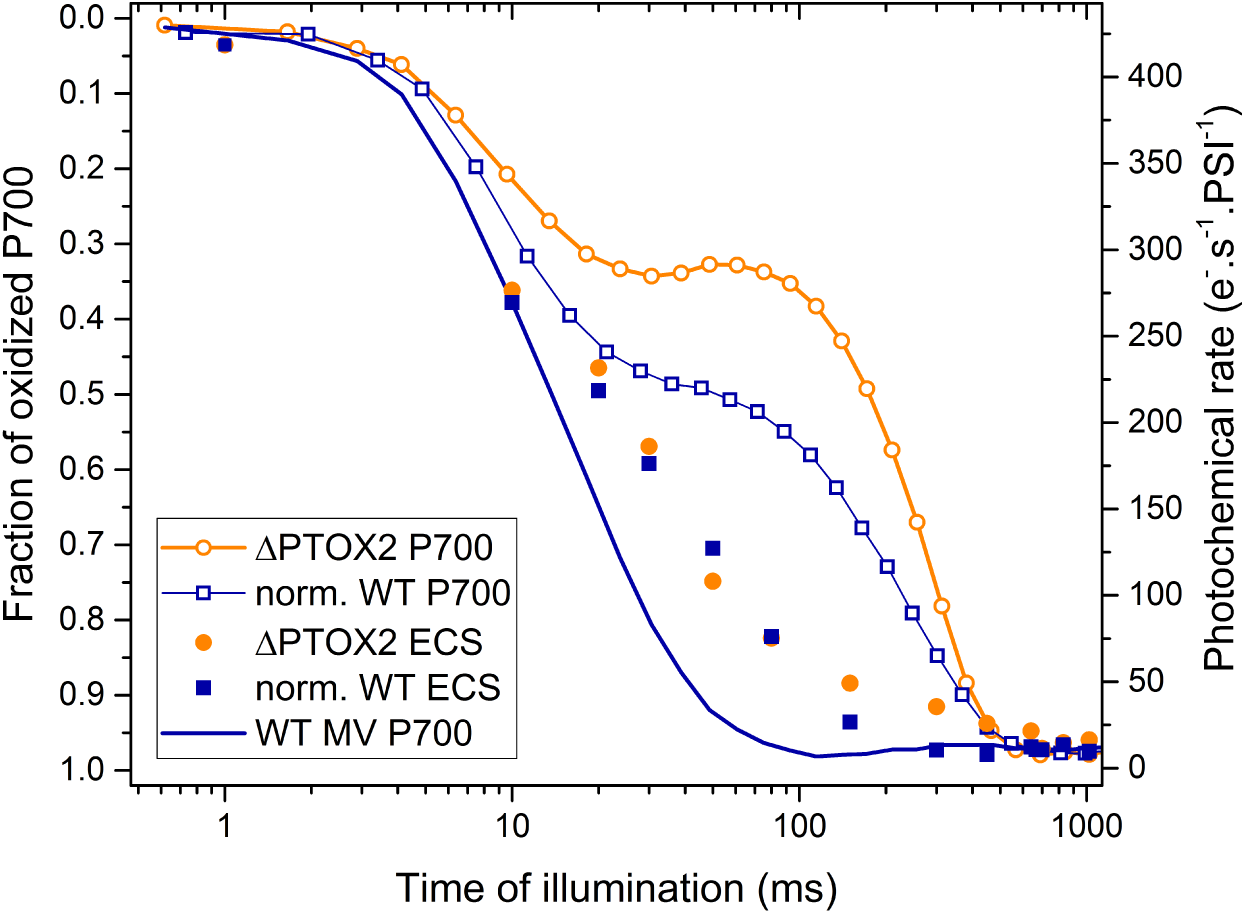
Initial PSI acceptor-side limitation smoothly cedes place to CEF upon dark-to-light transition

**Fig. S3, S4.**
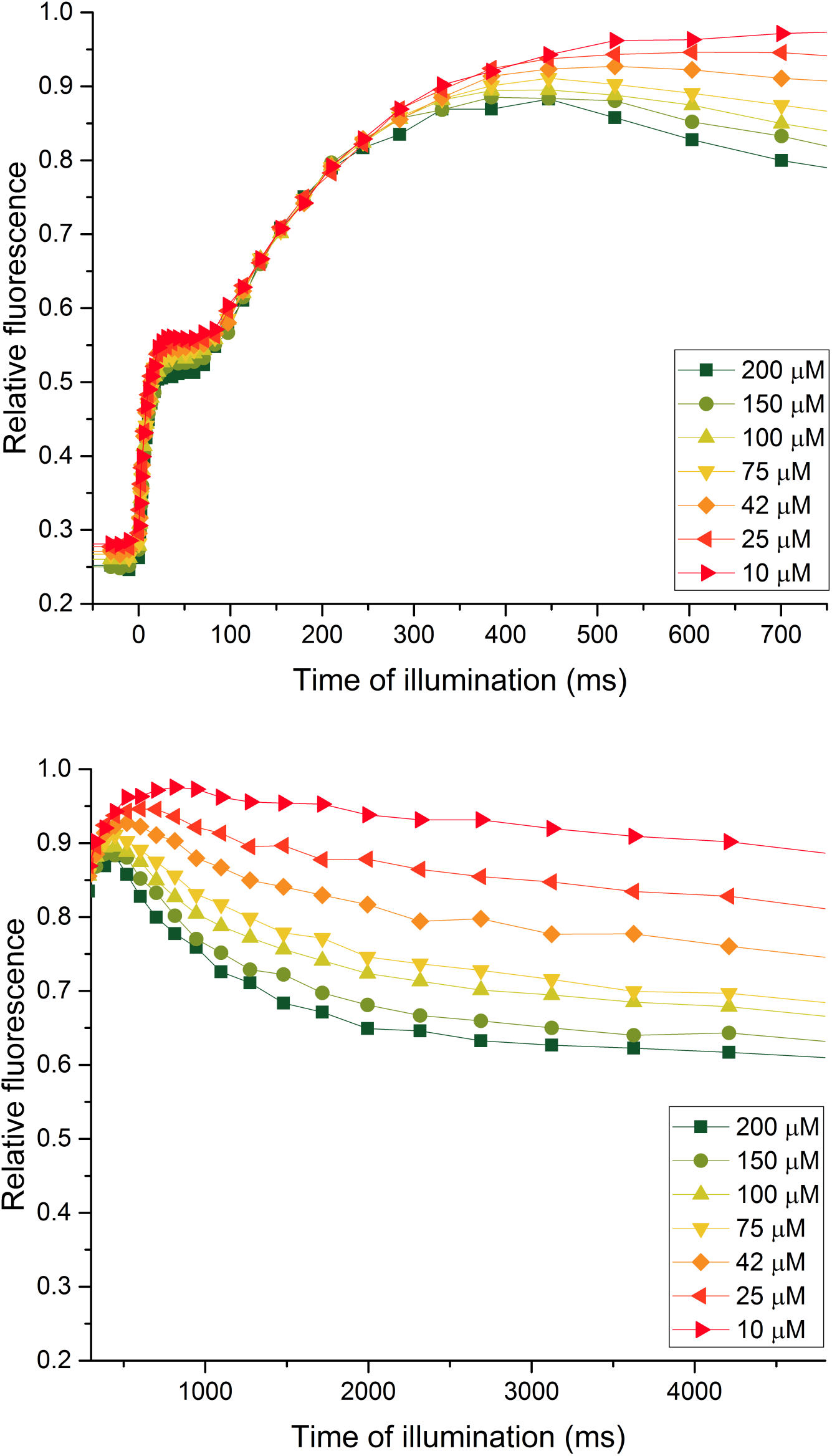
Fluorescence induction curves in a range of oxygen concentrations, first- and second phase

**Fig. S5.**
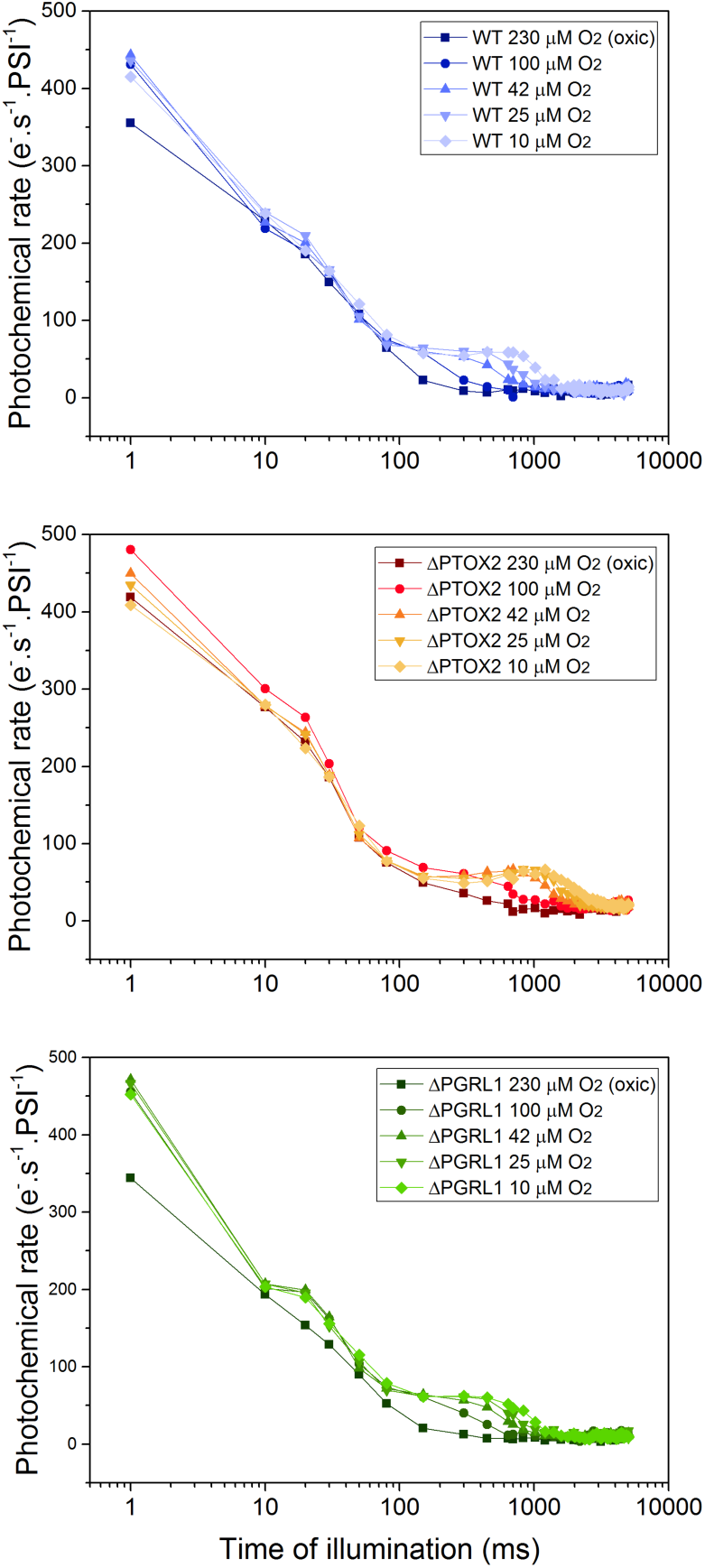
Full extent of the ECS measurements in hypoxia

